# Biogeography and Edaphic Factors Structure Coastal Sediment Microbial Communities More than Vegetation Removal by Sudden Vegetation Dieback

**DOI:** 10.1101/405464

**Authors:** Courtney Gorham, Aidan Barry, Beth A. Lawrence, Blaire Stevena

## Abstract

Development of sudden vegetation dieback (SVD), a phenomenon that causes the rapid mortality of salt marsh plants, specifically *Spartina alterniflora*, has affected large-scale alterations in Atlantic coastal systems, through the often-complete removal of vegetation. In this study, two wetlands that differ in the time since development of SVD were compared in order to study biogeographic and temporal patterns that structure coastal wetland microbial communities and their response to disturbance.

Biogeographic and edaphic factors that distinguished the two wetlands, such as differing salinity, water content, and soil carbon and nitrogen between the sites were more strongly associated with sediment microbial community structure than either sampling date or SVD development. In fact, no OTUs differed in abundance due to the season samples were collected, or vegetation loss due to SVD. This is not to say that SVD did not alter the composition of the microbial communities. The taxonomic composition of sediment communities in SVD-affected sediments was more heterogeneous between samples and a small number of OTUs were enriched in the vegetated sediments. Yet, these data suggest that coastal wetland sediment communities are predominantly shaped by environmental conditions and are generally resilient to temporal cycles or ecosystem disturbances.

**Importance:** One of the challenges of microbial ecology is predicting how microbial communities will respond to ecosystem change. Yet, few studies have addressed whether microbial responses to disturbance are consistent over space or time. In this study we employ SVD as a natural vegetation removal experiment and compare the sediment microbial communities between two geographically separated wetlands (*ca* 125 km). In this manner, we uncover a hierarchical structuring of the microbial communities, being predominantly governed by biogeography, with lesser effects due to disturbance, or temporal dynamics.

## Introduction

Coastal ecosystems are among the most productive on earth and have the potential to sequester and store carbon at rates of up to 50 times higher than other terrestrial ecosystems (1). The potency of coastal ecosystems as carbon sinks is attributable to their high primary productivity, as they can produce 40% more plant biomass annually than the same area of forest (2, 3). This plant fixed carbon is eventually delivered to the wetland sediments through litter, root exudates, or plant mortality, eventually becoming the substrate for microbial metabolism (4, 5). A defining feature of wetlands is periods of water saturation, with flooded sediments rapidly become anoxic. Anaerobic degradation of organic matter happens relatively slowly, and the rate of organic inputs from the vegetation occurs at a greater rate than losses by microbial respiration, resulting in a net accumulation of carbon in wetland sediments (6, 7). This stored carbon in coastal wetlands has been referred to as “blue carbon” and is an important component to mitigating the atmospheric carbon concentrations that are driving climate change (1, 8–10).

Coastal wetlands along the eastern coast of North America are experiencing sudden vegetation dieback (SVD), a phenomenon affecting low elevation salt marshes dominated by smooth cordgrass (*Spartina alterniflora*). SVD presents as an initial browning and thinning of the vegetation, with plant mortality occurring in periods as short as weeks (11, 12). Propagative rhizomes are killed by SVD, limiting plant regrowth and resulting in unvegetated patches that may remain for decades (11). Patches of SVD have been found to range from 300 m^2^ to 5 km^2^ with 50 to 100% plant mortality (13). The etiology of SVD remains controversial; fungal pathogens (14, 15), invasive crabs (16, 17), and drought (17) have been proposed as the cause, and likely interact or at least contribute to the development of SVD. Regional differences or complex interactions between factors may play a role in the difficulty in identifying a unified explanation of SVD development (18). The lack of an explanatory model for SVD development has driven most research to address the causes of SVD rather than the consequences of the loss of vegetation on the functioning of wetland ecosystems.

We previously documented that sediment microbial communities differed between SVD-affected sediments and stands of healthy *S. alterniflora* at a coastal marsh in Connecticut, U.S.A (19). SVD-affected sediments harbored reduced populations of bacteria in the phylum *Bacteroidetes,* whereas populations of sulfur-reducing bacteria (predominantly within the genus *Desulfobulbus*) were enriched in the SVD-affected sediments. Additionally, the SVD-affected patches supported 64% reduced CO_2_ emissions compared to healthy vegetated controls (19). Taken together, these observations indicate that SVD resulted in alterations in both the structure and function of sediment microbial communities. Yet, there is little known regarding the spatial or temporal scales at which the shifts in the microbial communities occur, or whether alterations in sediment microbial communities are similar between geographically isolated wetlands.

In the present study, we compared sediment microbial communities between two salt marshes both experiencing current outbreaks of SVD. However, the time since SVD development differed between the two sites (5 versus. 10 years). To examine the relative role of vegetation, we examined sediment microbial communities in summer (July) during peak plant activity, and in fall (October) when salt marsh plants begin senescence. We predicted that the microbial communities would differ between the two field sites due to biogeography, but would show similar responses to SVD, such that the sites experiencing SVD would potentially be more similar to each other than they were to healthy locations at the same field site. We further expected that the sediment microbial communities response to SVD would be muted in fall sampling, when plants in the vegetated plots were not active and producing root exudates for the microbial populations. In this manner, alterations in the coastal wetland sediment communities could be linked to spatial biogeography, and to temporal dynamics related to the time from disturbance, and seasonal plant activity.

## Results

### Site descriptions

Sediment samples were collected in 2015 from Hammonasset Beach State Park (hereafter referred to as Hammonasset) on July 22^nd^ and October 8^th^ of 2015 and at Narragansett Bay National Estuarine Research Reserve (hereafter referred to as Narragansett) on July and October 13^th^. The two sites are separated by *ca.* 125 km (Fig. 1A). All sampling was performed at low tide, as this is the only period where SVD sediments are exposed for collection (Fig. 1B).

**Figure 1.**
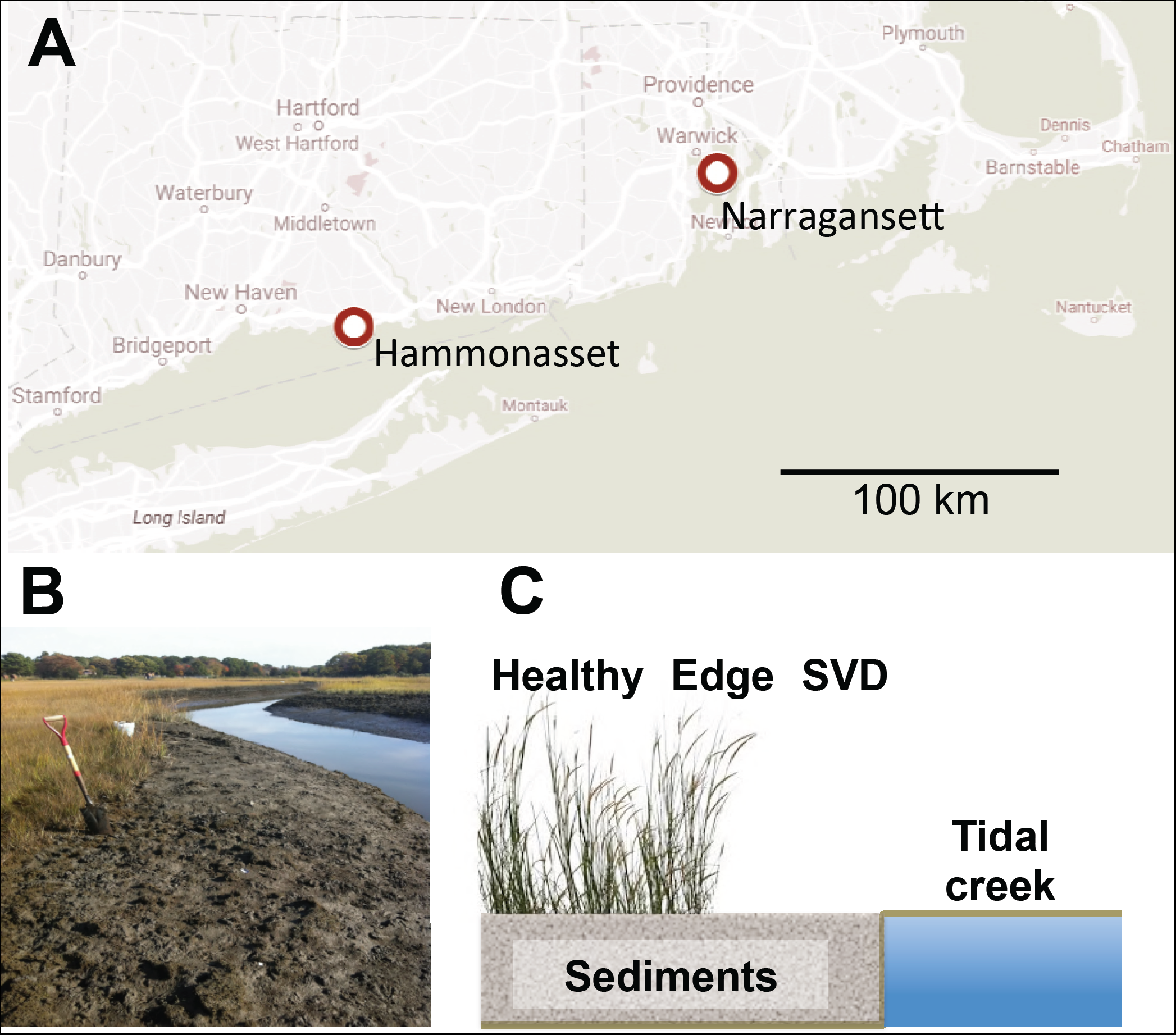
A). Map of the East coast of the USA showing sampling sites. The sites were approximately 125 km apart B). Photograph of a site experiencing SVD at Hammonasset Beach State Park. B). Schematic diagram displaying the sampling strategy. Samples consisted of a transect originating at the SVD sites, which occur adjacent to the tidal creeks. Edge samples were collected from sediments in the region where vegetation began to grow, and healthy sediments were collected from sites within stands of flourishing *S. alterniflora* approximately 1 meter distant from the edge sample.

The dominant vegetation at both sites is *S. alterniflora* and both sites have unvegetated patches due to SVD. At Hammonasset, outbreaks of SVD were first documented in 1999 and many patches have remained unvegetated since. At Narragansett, SVD was first reported in 2010 and represents a more recent occurrence of SVD. At each site, we established three transects perpendicular to tidal creeks, and sampled sediments at three locations (Fig. 1B,C): in SVD patches adjacent to tidal creeks (“SVD”), in the transitional edge where vegetation was first encountered (“Edge”), and in healthy, well-established stands of *S. alterniflora ca.* 1 m from the edge sample (“Healthy”; Fig. 1C). Thus, samples differ in both their vegetation status and in their spatial relationship to the local tidal creeks. Together, we collected 18 sediment samples (2 sites x 3 transects x 3 locations) during each of the summer and fall sampling campaigns.

### Sediment chemistry

The two sites differed significantly (P < 0.05) in all sediment chemistry variables except pH (F_1,31_ = 3.9, P = 0.056). In general, Narragansett had higher soil electrical conductivity (EC; F_1,31_ = 6.3, P= 0.017), soil moisture (F_1,31_ = 141.7, P < 0.001), soil %C (F_1,31_ = 199.9, P < 0.001) and %N (F_1,31_ = 292.3, P < 0.001), but a lower soil C:N than Hammonasset (F_1,24_ = 38.0, P < 0.001; Table 1). None of the measured sediment variables differed among vegetation zones nor season (P > 0.05). However, we observed an interaction between site and season of sampling in C:N ratios (F_1,24_ = 8.5, P= 0.007), where the Narragansett sediments had greater C:N during the summer sampling than the Hammonasset wetland, but had greater C:N when sampled in the fall.

**Table 1.**
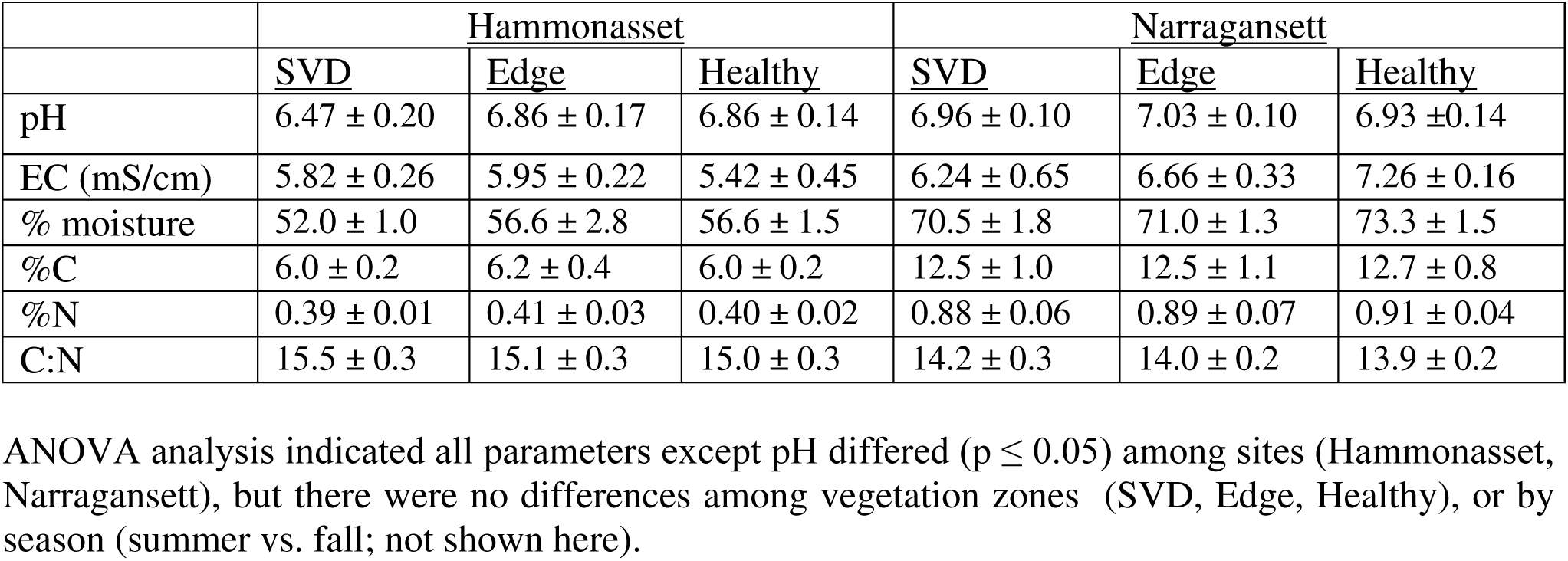
Sediment characteristics (mean ± SE) of two northeastern (USA) coastal wetlands affected by SVD.

|  | <u>Hammonasset</u> |  |  | <u>Narragansett</u> |  |  |
| --- | --- | --- | --- | --- | --- | --- |
|  | <u>SVD</u> | <u>Edge</u> | <u>Healthy</u> | <u>SVD</u> | <u>Edge</u> | <u>Healthy</u> |
| pH | 6.47 ± 0.20 | 6.86 ± 0.17 | 6.86 ± 0.14 | 6.96 ± 0.10 | 7.03 ± 0.10 | 6.93 ± 0.14 |
| EC (mS/cm) | 5.82 ± 0.26 | 5.95 ± 0.22 | 5.42 ± 0.45 | 6.24 ± 0.65 | 6.66 ± 0.33 | 7.26 ± 0.16 |
| % moisture | 52.0 ± 1.0 | 56.6 ± 2.8 | 56.6 ± 1.5 | 70.5 ± 1.8 | 71.0 ± 1.3 | 73.3 ± 1.5 |
| %C | 6.0 ± 0.2 | 6.2 ± 0.4 | 6.0 ± 0.2 | 12.5 ± 1.0 | 12.5 ± 1.1 | 12.7 ± 0.8 |
| %N | 0.39 ± 0.01 | 0.41 ± 0.03 | 0.40 ± 0.02 | 0.88 ± 0.06 | 0.89 ± 0.07 | 0.91 ± 0.04 |
| C:N | 15.5 ± 0.3 | 15.1 ± 0.3 | 15.0 ± 0.3 | 14.2 ± 0.3 | 14.0 ± 0.2 | 13.9 ± 0.2 |

ANOVA analysis indicated all parameters except pH differed ( $p \leq 0.05$ ) among sites (Hammonasset, Narragansett), but there were no differences among vegetation zones (SVD, Edge, Healthy), or by season (summer vs. fall; not shown here).

### Relationship between sequence datasets

Constrained Analysis of Principal Coordinates (CAP) ordination was used to investigate patterns in the relationships between the sequence datasets. The samples from the two field sites were clearly distinguished (Fig. 2; PERMANOVA P=0.001), suggesting that the sediment microbial communities were significantly different between the two wetlands. Vegetation status was also associated with a significant difference in clustering of the datasets (PERMANOVA P=0.028). Furthermore, the interaction between the date of sampling and vegetation status was not significant (P=0.995), suggesting that the date of sampling did not influence the microbial community composition associated with the different vegetation conditions. Date of sampling was not significant factor in sample clustering (P = 0.20). Taken together, these data suggest that coastal sediment microbial community composition is primarily structured by the edaphic factors associated with biogeography, followed by vegetation removal by SVD, with a very small contribution of sampling date.

**Figure. 2.**
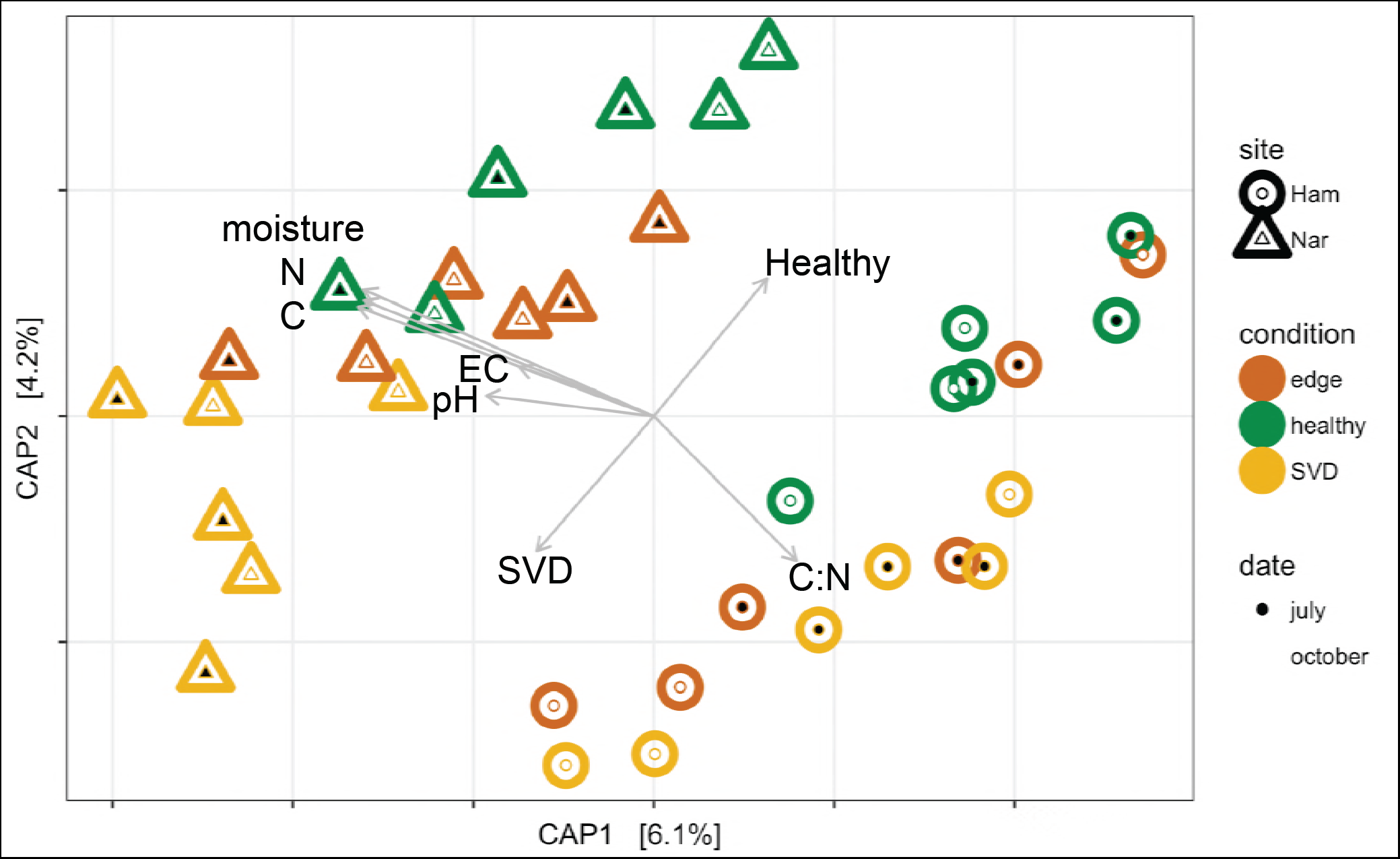
Canonical Analysis of Principal Coordinates (CAP) of OTU abundance data in the sequence datasets. Inter-sample distances were calculated with the Bray-Cutis metric using rarefied OTU count data. The percent variance explained by each of the CAP axes is indicated. Explanatory variables are indicated by arrows (data presented in Table 1).

### Microbial diversity

To measure alpha diversity, the datasets were rarified to the same number of sequences (7,689) and three diversity metrics were calculated, the number of observed OTUs, Shannon’s diversity index, and inverse Simpson’s index (Fig. 3). The average number of OTUs recovered from the Hammonasset samples was 4,594 (±521) and 4,636 (±378) for Narragansett. The Shannon’s diversity index for both Hammonasset and Narragansett samples was 7.95, and the inverse Simpson’s index was 983 for Hammonasset and 895 for Narragansett (Fig. 3). Furthermore, there was no apparent diversity pattern between samples collected during different seasons or from different vegetation zones. Together, these data indicate that sediment microbial diversity was not affected by biogeography, date of sampling, or vegetation status.

**Figure. 3.**
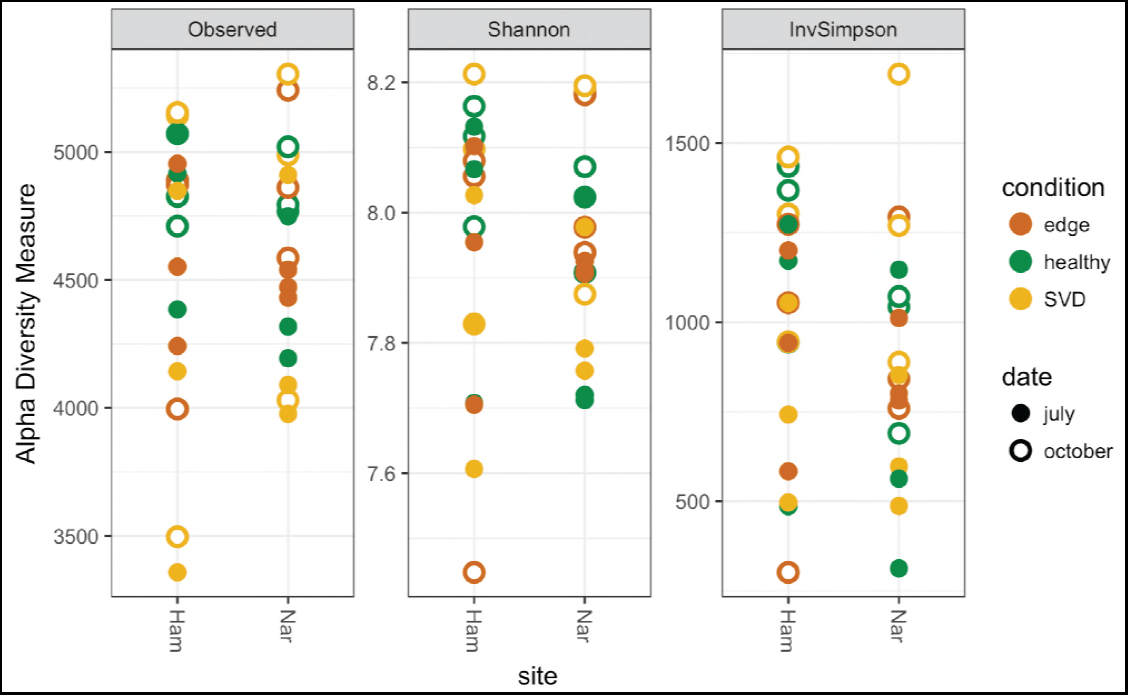
Alpha diversity of sequence datasets, separated by field site (Ham=Hammonasset, Nar=Narragansett). Three diversity indices were calculated using OTU abundance in the rarefied datasets. The number of observed OTUs (Observed), Shannon’s diversity index (Shannon), and the Inverse Simpson’s Index (InvSimpson). Each point represents the value from a single dataset.

### Taxonomic composition of datasets

Sequence reads were classified to the phylum level to compare the composition of the bacterial communities between the sites (Fig. 4). In general, phyla were present at the two sites in similar proportions. For example, at both sites the two dominant identified phyla were Proteobacteria and Bacteroidetes (Fig. 4). Both sites also harbored a relatively large proportion unclassified bacterial sequences, suggesting a large fraction of uncharacterized bacterial diversity in the sequence datasets.

**Figure. 4.**
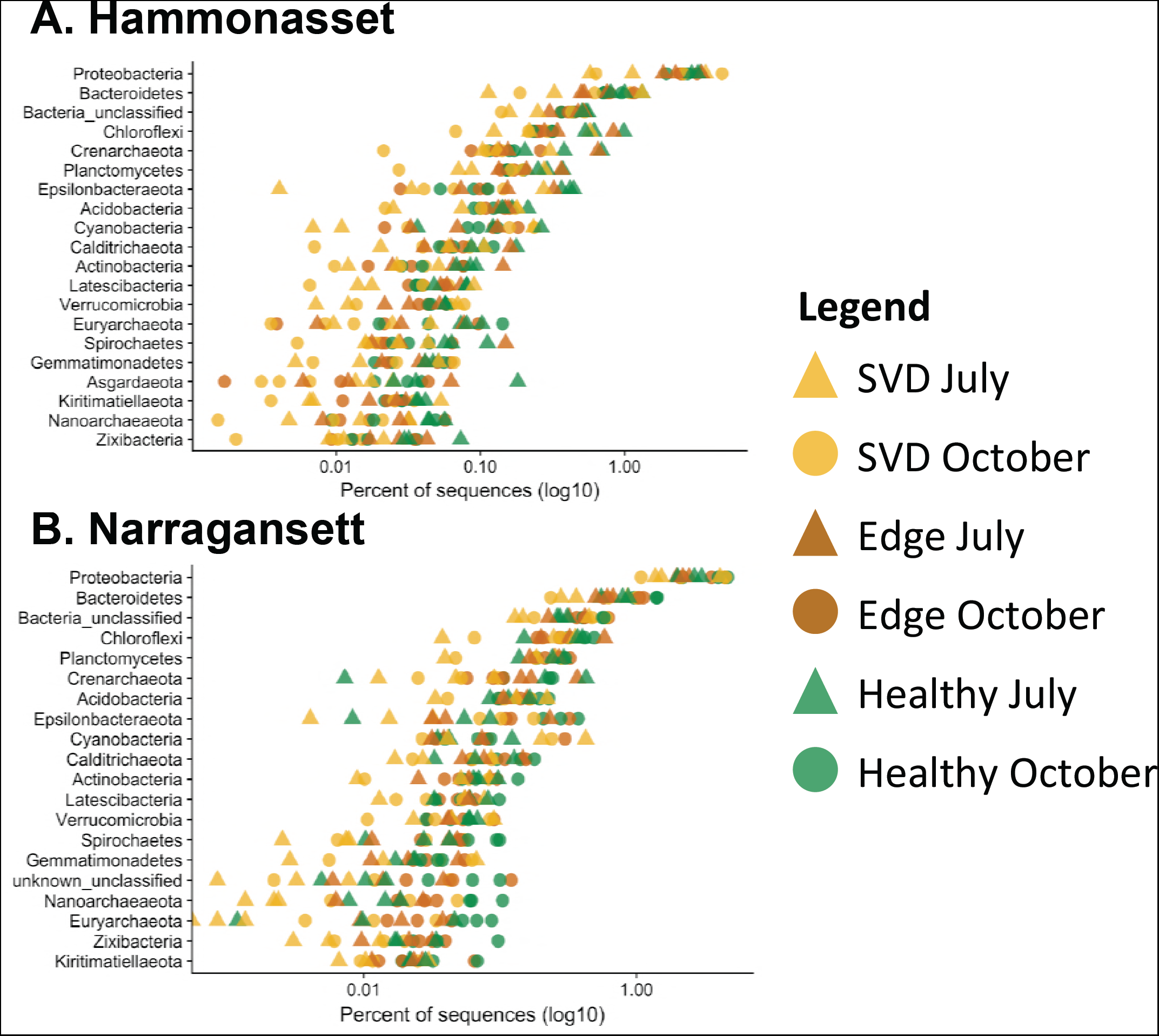
Phylum level taxonomic bins in the sequence datasets, separated by field site. Each point is the average of three replicate samples. Only the 20 most numerically abundant phlyla (sorted by abundance across all samples for each site) are displayed.

A common observation across the datasets was that samples with phyla that showed large deviation from the mean were predominantly from the SVD conditions. Yet, these shifts were not consistent across replicate samples or with sampling date (Fig. 4). In this regard, no phylum level taxonomic bins were found to be significantly different in relative abundance when tested for either date of sampling or vegetation status. Thus, these data suggest a part of the sediment microbial community response to SVD is to increase the taxonomic heterogeneity between samples, rather than a consistent shift of specific taxonomic ranks.

### Differentially abundant OTUs due to site

A total of 23 OTU’s (97% sequence identity) were identified as significantly different in relative abundance due to site, 9 significantly enriched at Hammonasset and 14 significantly enriched at Narragansett (Fig. 5). The differentially abundant OTUs belonged to five phyla and could be further classified to 12 taxonomic ranks representing the deepest level to which the OTUs could be reliably assigned (Table S1). There was no obvious pattern in the taxonomy of the differentially abundant OTUs. In fact, several OTUs were identified to taxa that were significantly more abundant in both of the field sites. For instance, two OTUs identified as significantly more abundant at Hammonasset were classified to the genus *Calothrix* along with one of the OTUs that was enriched at Narragansett (Table S1). In this respect, these data suggest that at least a portion of the differentially abundant OTUs due to site may represent functionally redundant species adapted to the local edaphic factors.

**Figure. 5.**
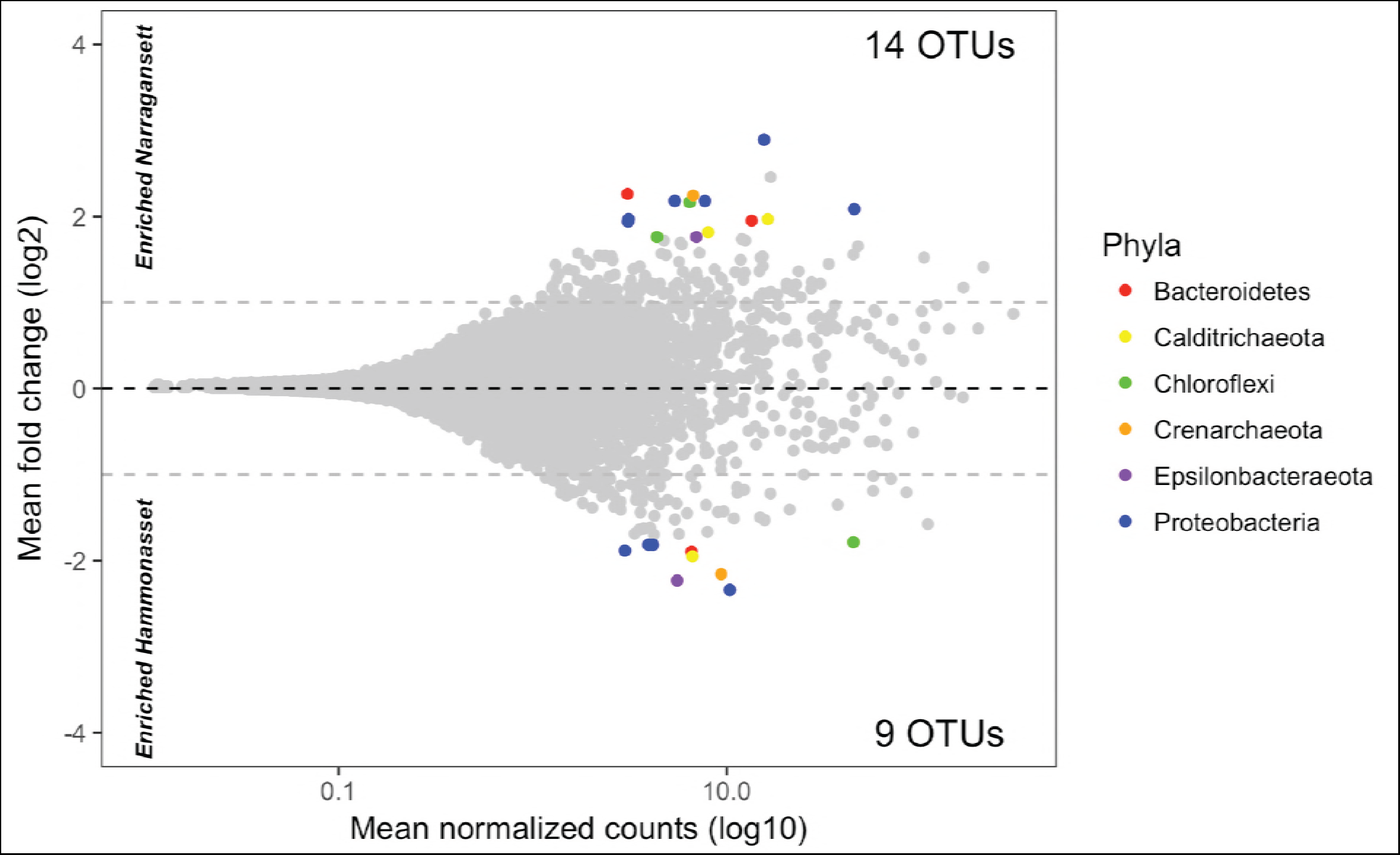
Mean OTU counts versus mean fold change in abundance for each OTU in the dataset. Points represent the mean values for 36 samples. OTUs with a significant difference in abundance are colored by the phylum to which they were classified. The dashed lines indicate a two-fold difference in relative abundance between sites. The number of OTUs enriched at each site is indicated in the panel. A full list of the differentially abundant OTUs is presented in Table S1.

### Differentially abundant OTUs due to sampling date

Samples were collected in July and October to test for temporal dynamics in the sediment communities. Overall, the majority of OTUs did not show a large change in relative abundance, rarely surpassing a two-fold difference between sampling dates (Fig. 6). None of the OTUs were identified as significantly different in abundance. Thus, these data suggest that sampling date was a small factor in driving sediment community structure. This further matches the ordination results in which sampling date was not a significant factor in sample clustering (Fig. 2).

**Figure. 6.**
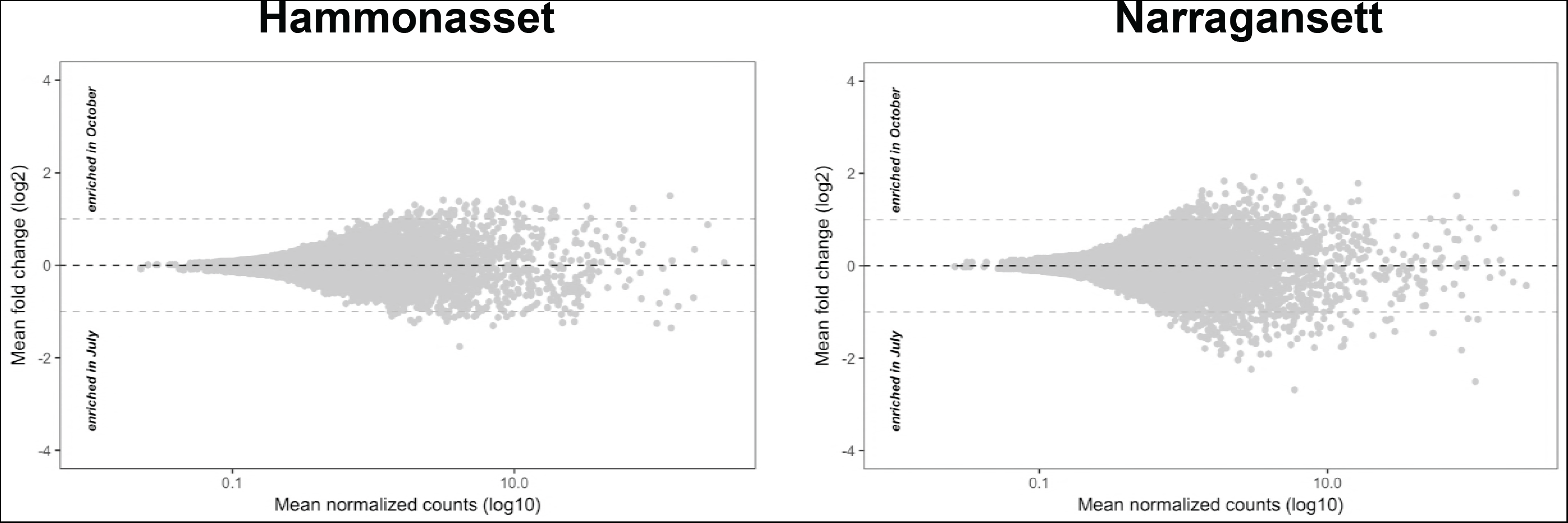
Mean OTU counts versus mean fold change in abundance for each OTU in the dataset grouped by site. Points represent the mean values for 18 samples. No OTUs were found to be significantly different in abundance. The dashed lines indicate a two-fold diffierence in relative abundance between sample dates.

### Differentially abundant OTUs due to vegetation status

OTU abundance among the samples differing in vegetation status was investigated (Fig. 7). Large portions of the most abundant OTUs in the datasets were present in roughly equal abundance between all three vegetation conditions (inner triangle Fig. 7). Additionally, no OTUs were identified as significantly different due to vegetation status at either site (ellipses Figure 7). Yet, there was a clear trend of certain OTUs being more abundant in the vegetated sites (both healthy and edge), with few OTUs showing enrichment in the SVD sediments. Thus, these data suggest that there are certain OTUs that trend toward being more abundant in the vegetated samples even if they did not rise to the level of significance. When taxonomy was mapped onto the OTUs, there was no readily apparent pattern in the OTUs that were more abundant in the vegetated sites as they were represented by multiple phyla (Fig. 7).

**Figure. 7.**
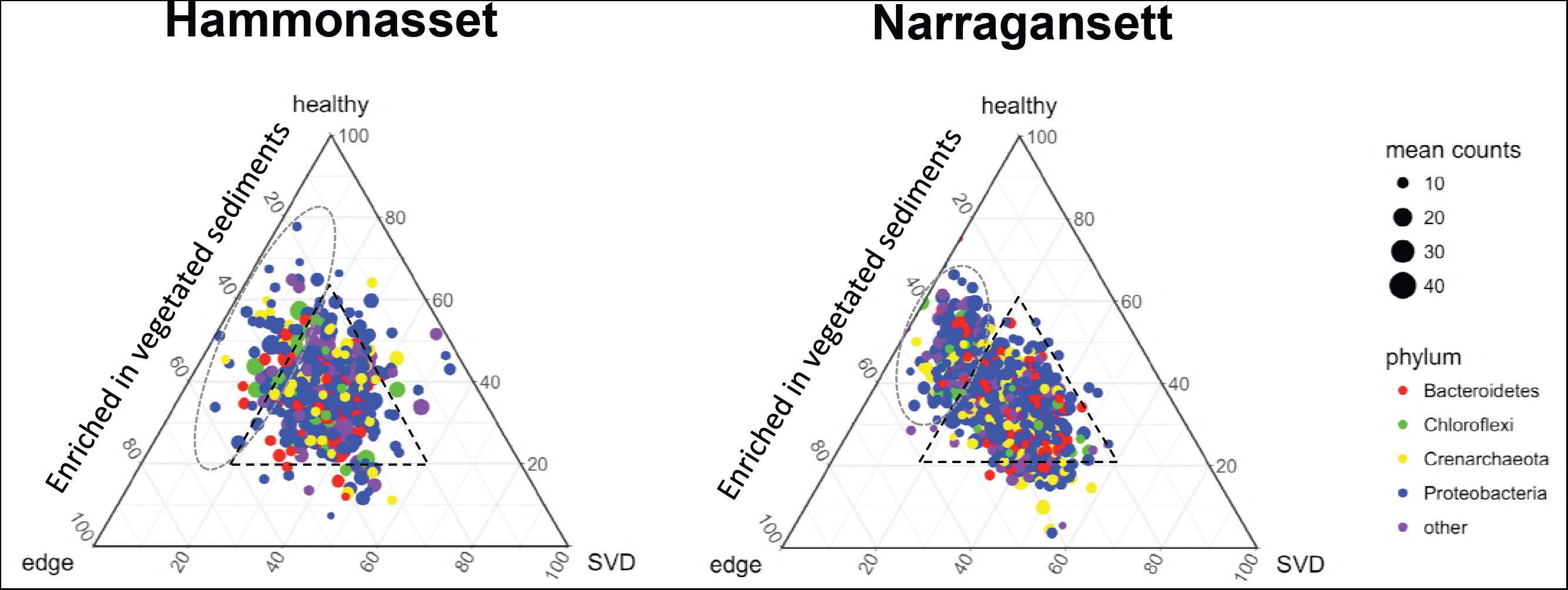
Ternary diagrams displaying relative abundance of OTUs among the three vegetation conditions. The 500 most abundant OTUs displayed and colored based on the phylum to which they were classified (the four most common phyla are indicated, with remainder assigned to “other”). The size of the point indicates the mean count of each OTU across the datasets (n=18). Only OTUs with at a sequence count of at least 10 were retained to examine OTUs that could potentially be shared between all three conditions. The central triangle denotes the centered mean of the diagram and thus OTUs present in each condition in roughly equal relative abundance. The ellipses denote OTUs more abundant in vegetated samples (edge and healthy) and is a visual aid only. Note that no OTUs were significantly diffierent in abundance due to vegetation condition.

## Discussion

The results of this study demonstrate edaphic factors related to geography were a larger diver of sediment community composition than the date of sampling or vegetation removal by SVD. Sediments from Narragansett had higher soil moisture, greater electrical conductivity, and higher C and N content, indicating that tidal waters may have more frequently inundated the Narragansett sites (Table 1). While sediments were collected systematically along perpendicular transects from tidal creeks at both sites, it is possible that Narragansett SVD patches occurred lower in the tidal frame and thus were wetter, saltier, and enriched in organic matter, key factors driving microbial community composition.

Previous studies have similarly found that geography is a large driver of bacterial community composition. Regional differences in Louisiana salt marshes were at least as large of a predictor of bacterial community composition as those between the rhizosphere of different plants (*S. alterniflora* and *Juncus roemerianus;*(20)). Similarly, ammonia-oxidizing communities (bacteria and archaea) showed larger differences between regions associated with soil moisture and nitrogen content than due to contamination during the Deepwater Horizon oil spill (21). Despite biogeography being a large influence on sediment microbial communities, the structure of the sediment microbial communities was similar between the two field sites, being composed of the same dominant taxa (Fig. 4), harboring similar levels of microbial diversity (Fig. 3), and only a relatively small number of OTUs being identified as significantly different in relative abundance between the sites (Fig. 5). A subset of the differentially abundant OTUs belonged to taxonomic groups specifically enriched in a certain field site. For example, an OTU identified to the genus *Mariprofundus* was enriched at Hammonasset (Table S1). These organisms have been associated with a role in iron oxidation in marine systems (22), and may point to differences in iron cycling between the sites. In contrast, several OTUs belonged to taxonomic ranks identified as more abundant in both field sites (Table S2). This could indicate that the differentially abundant taxa are largely functionally redundant but have adapted to the different biogeographic and edaphic factors that differentiate the two wetlands.

Coastal wetlands are temporally dynamic systems. Diurnal cycles in the tidal cycle, light, and photosynthesis rates drive changes in the magnitude of sediment respiration (23). Up to 76% of the detectable methane emissions from coastal wetland sediments are released during tidal immersion (24). Finally, seasonal patterns result in large alterations in a multitude of environmental factors, including temperature and plant physiology. Carbon flux from sediments is generally lower in the non-growing seasons, even when accounting for lower average temperatures (25) and plant activity in summer may regulate biogeochemical processes such as iron cycling (26). Thus, these data support that coastal sediment microbial activity is largely driven by biotic and abiotic factors that vary at a variety of time scales. However, studies characterizing temporal patterns in microbial community assembly are notably sparse (27–29). We collected samples in July and October to investigate if the sediment microbial communities showed significant changes in composition related to season. Date of sampling was insignificant as a factor clustering the sequence datasets (Fig. 2) and the OTUs in the datasets were present in similar abundances at both time points with no OTUs being identified as significantly different in abundance due to sample date (Fig. 6). Taken together, these data suggest that the sediment microbial communities were largely similar in summer and fall samples, suggesting a limited role for seasonal dynamics in shaping the sediment communities. The samples were limited to a two-point time course therefore a finer-grained analysis may be required to disentangle a more nuanced response of these communities to temporal cycles. Yet these observations suggest that while microbial activity is responsive to the temporal dynamics in coastal wetlands, alterations in activity may be a poor predictor of community composition.

Finally, we employed the development of SVD as a natural experiment to assess the impact of an ecosystem disturbance on the sediment communities. At a landscape scale, the complete loss of vegetation appears to be a dramatic disturbance that would presumably translate into similarly large shifts in the sediment microbial communities. We previously showed that sites at Hammonasset experiencing SVD supported significantly lower populations of bacteria within the phylum *Bacteroidetes* and an elevated relative abundance of sulfate reducing bacteria (19). Yet, in this study no OTUs were identified as significantly different in relative abundance. The lack of significant differences could be due to the relatively low replication per individual sample date and site, the depth of sequencing, or the added variability of identifying differences between samples collected on different dates. It is important to note, that while this study did not identify any OTUs significantly altered in abundance due to SVD, that does not indicate that there was no effect on the sediment communities. For example, CAP analysis identified a significant difference in the clustering of samples under different vegetation statuses (Fig. 2), the taxonomic makeup of the sediment communities at the phylum level was more heterogeneous in the SVD samples (Fig. 4), and there was a clear trend in OTUs that were enriched in the vegetated sites (edge and healthy) compared to SVD sites (Fig. 7). These data indicate that to the extent that there are shifts in the microbial communities due to SVD, they are likely limited to relatively rare community members and do not involve large shifts in relative abundance. The practical relevance of these observations is that the sediment microbial communities are also likely to respond well to any restoration efforts. For example, we previously demonstrated that SVD-affected sediments are capable of maintaining *S. alterniflora* germination and growth in greenhouse experiments (19). In this respect, we propose that restoration efforts for SVD-affected sites should primarily focus on the plant communities, as any alterations in the sediment microbial communities do not appear to have any deleterious effects on plant health.

A multitude of studies have similarly found that disturbance can lead to relatively small shifts in community composition of salt marsh sediment communities. For instance, increasing nitrogen loading to wetland sediments caused decreases in the metabolically active microbial populations without a concurrent alteration in the total community composition (30) or may be generally limited to specific nitrogen cycling populations (31). Even reciprocal transplants between salt marshes resulted in negligible changes in sediment microbial communities (32). A metagenomic survey of two tidal creeks, one of which had received more than 40 years of sewage effluent from its headwaters, found little difference in the taxonomic profile of sediment communities but significant differences in the abundance of nitrogen cycling genes, suggesting shifts in the functional potential of the community without concurrent shifts in community membership (33). In this regard, sediment microbial communities have proven themselves to be resilient to disturbance.

## Conclusion

Microbial communities are responsible for many of the ecosystem functions of coastal wetlands, particularly those for carbon sequestration and storage. Thus, characterizing the resiliency of these communities to disturbance and their natural variation with biogeography and time will be central to modeling their activities, and future in a changing environment. In this study, we show that the sediment microbial communities were relatively unaffected by an ecosystem disturbance, vegetation removal by SVD, at two different wetlands. Furthermore, temporal patterns in the community were small with little change in the composition of the communities between summer and fall. This suggests that the taxonomic makeup of the sediment microbial community was relatively stable in the face of seasonal dynamics and disturbance. These data support that the taxonomic makeup of sediment microbial communities are largely resilient to diverse environmental perturbations and future work may need to focus more specifically on the metabolic activities of the microbial populations.

## Materials and Methods

### Field sampling

Sediment samples consisted of *ca.* 5g of material collected with an ethanol-sterilized spatula. Sediments were collected from the upper 1 cm to focus on surface communities. The sediment samples were transferred to Whirl-Pak™ bags, placed on dry ice in a cooler, and transported to the Connecticut Agricultural Experiment Station (New Haven, CT) where they were stored at −80°C until DNA extraction.

### Sediment chemistry

Sediment samples were stored frozen (−18°C) and thawed prior to analysis. Soils were sieved through a 2-mm mesh screen to remove belowground biomass and subsamples were analyzed for several sediment parameters. Soil pH and electrical conductivity were estimated on 10g subsamples diluted with 50 mL of deionized water and quantified using Orion Star A215 pH Conductivity Meter Orion with Ross Ultra pH/ATC Triode (8157BNUMD) and Orion Conductivity Cell (013005MD) probes. We dried subsamples at 105°C for >24hours and then weighed to estimate soil moisture. Subsamples were also pulverized in a ball-mill, rolled in tins, and analyzed for %C and %N (Costech ECS 4010 CN Analyzer). Every ten samples we ran analytical triplicates to examine sample heterogeneity and observed <20% standard deviation for all soil parameters.

### DNA extraction, 16S rRNA gene amplification, and sequencing

Samples were processed as described previously (19, 34). Briefly, total environmental DNA was extracted from sediments using the DNeasy PowerSoil Kit (Qiagen), using standard protocols. DNA extractions were verified by gel electrophoresis. 16S rRNA genes were amplified with the universal primers 515F and 806R (35), which also included Illumina adaptor sequence. Cycling conditions were as follows: Initial denaturation 95°C for 3 min; 35 cycles of 95°C for 45 s, 55°C for 60 s, 72°C for 60 s; a final extension at 72°C for 10 min. PCR amplification was checked by gel electrophoresis, verifying a *ca.* 300 b.p. amplification product. PCR products were purified using the QIAquick PCR purification kit (Qiagen). Sequence indexing was performed using the index PCR procedure, employing the Nextera DNA Library Prep Kit (Illumina Inc., San Diego, CA). Following indexing, PCR amplicons were purified as above and pooled in equal molar concentrations (1 µg). The resulting DNA was sequenced on the Illumina HiSeq platform at the Yale Center for Genome Analysis with standard protocols on the HiSeq2500 employing paired end 2 × 150 chemistry.

### Sequence processing

Paired end sequences were assembled into contigs using the make.contigs command with default parameters in the mothur (36) software package, only retaining contigs of at least 291 bases in length. Each contig was further screened to remove any sequences with any ambiguous nucleotide calls or homopolymers of ≥7 bases. Potentially chimeric sequences were identified with the mothur implementation of VSEARCH (37) and removed from the dataset. Sequences were clustered into operational taxonomic units (OTUs) with the OptiClust algorithm in mothur (38). For analyses of diversity and composition, an OTU definition of ≥97% sequence identity was employed. Taxonomic assignment of reads was performed with the mothur implementation of the Naïve Bayesian Classifier (39) against the SILVA (40) ribosomal gene database as maintained by mothur. Sequences with confidence scores of ≥80% were considered to be reliably classified.

### Statistical analyses

For each of the 6 sediment variables (pH, EC, soil moisture, soil %N, soil %C, soil C:N), we tested for normality with Shapiro-Wilks tests and log-transformed if necessary. We used a 3-way, interactive ANOVA (site * vegetation type * season); if there were no significant interactions (true in all cases, except soil C:N), we simplified to an additive ANOVA (site + type + season).

OTU abundance data was uploaded to the phyloseq R software package (41, 42) for calculation of ordination plots and calculations of alpha diversity. CAP analysis was performed on data randomly rarified to the sample size of the smallest sequence dataset (7,689 sequences). Inter-sample distances were calculated with the Bray-Curtis metric and PERMANOVA statistics were calculated with the Adonis function of the vegan R package (43). For testing for significant clustering of date and vegetation status samples the model included an interaction term to test if there was a significant interaction between the variables.

Differentially abundant OTUs were identified by calculating the log_2_-fold ratio using the negative binomial generalized linear framework of the DESeq2 software package (44). Unnormalized OTU count data was used for all tests. P-values were adjusted for multiple tests with a Benjamini–Hochberg false discovery rate correction, and a threshold P-value of 0.01 was used to prevent the likelihood of false positives. We used the log_2_–fold ratio of the relative abundance and mean normalized counts to display differentially abundant OTUs in a Bland–Altman plot. The model for detecting differentially abundant OTUs accounted for the nested nature of the sampling as site + date + condition (vegetation status).

### Sequence availability

All sequences generated in this study are available in the NCBI sequence read archive under the BioProject ID: PRJNA488460.

## Acknowledgements

The authors would like to acknowledge Kenny Raposa and Henry Alves for field site access. We would also like to thank Dr. Wade Elmer and Peter Thiel for assistance in field sampling. This work was supported by the USDA National Institute of Food and Agriculture, Hatch project 1006211.

